# Genetic drift decouples Fisherian trait-preference coevolution from genetic correlations in finite populations

**DOI:** 10.64898/2026.09.10.750770

**Authors:** Kuangyi Xu

## Abstract

A central mechanism of sexual selection theory is Fisher’s process, which refers to the coevolution of male traits and female preferences, in which preference is indirectly selected through genetic association with male trait alleles built up via mate choice. However, empirical studies often fail to detect strong trait-preference genetic correlations, raising doubts about the importance of Fisherian selection in nature. Notably, the theoretical expectation that genetic correlations are essential for trait-preference coevolution through Fisherian selection derives largely from models assuming infinitely large populations, whereas real populations are finite. Using population genetic models, I show that interactions between Fisherian selection and genetic drift can fundamentally decouple trait-preference genetic correlations from the evolution of male traits and female preferences. Genetic drift generally reduces expected trait-preference correlations but simultaneously promotes the expected increase in female preference frequency beyond deterministic predictions. Consequently, in populations of realistic sizes, trait-preference correlations may often be weak or even negative, particularly when recombination among trait and preference loci is infrequent, but substantial trait-preference coevolution can still occur. Therefore, the strength of genetic correlations may be an unreliable indicator of the extent of trait and preference coevolution, offering a potential resolution to a longstanding dilemma in sexual selection theory.

## Introduction

The prevalence of exaggerated display traits in males and female preference for these seemingly non-adaptive traits have received long-standing interest in evolutionary biology. Darwin (1871) proposed that elaborate display traits may evolve through higher mating success when females preferentially mate with males bearing such traits. Fisher (1930) further argued that female preferences can evolve through indirect selection arising from genetic correlations with preferred male traits, as mating between choosy females and preferred males yields offspring inheriting both preference and display alleles. The evolution of preference in turn strengthens selection on the male trait, generating self-reinforcing feedback that can drive both traits and preferences toward increasingly extreme values (Lande 1981; Kirkpatrick 1982). This trait-preference coevolutionary process—in which selection on female preferences is indirect through its genetic association with male traits—is known as Fisherian sexual selection, and has since become a central mechanism in sexual selection theory (Mead & Arnold 2004; Prum 2010; Kuijper et al. 2012).

Measuring trait-preference correlations has been a key test of the operation of Fisherian selection in nature, because the strength of the correlation determines whether exaggeration of male traits and female preferences can occur (Lande 1981; Hall et al. 2000; Xu et al. 2023), and whether indirect selection on preferences can overcome potential direct costs (Bulmer 1989; Kirkpatrick & Barton 1997; Fry 2022; Servedio 2025). Nevertheless, empirical studies have found that such correlations are generally not significantly different from 0, even in populations with substantial genetic variation in traits and preferences (Ritchie et al. 2005; Zhou et al. 2011; Greenfield et al. 2014). Several explanations have been proposed to reconcile this dilemma. One explanation is that traits and preferences can evolve via alternative processes that do not require strong genetic correlations, such as context-dependent female preference shaped by social experience and encountered male phenotypes (Bailey & Moore 2012). A second explanation posits that genetic correlations between traits and preferences may build up only intermittently, due to interruptions by immigration and/or spatiotemporal environmental heterogeneity, especially when male traits and preferences are plastic (Greenfield & Rodriguez 2004; Kokko & Heubel 2008; Chaine & Lyon 2008; Greenfield et al. 2014).

However, whether trait-preference genetic correlations provide a reliable indicator of the extent of trait-preference coevolution through Fisherian selection in nature requires careful scrutiny. Natural populations are finite, in which case interactions between selection and genetic drift may fundamentally alter evolutionary dynamics. Population genetic theory has shown that in finite populations, two independent beneficial alleles develop negative associations on average, thereby impeding each other’s fixation, a phenomenon known as the Hill-Robertson effect (Hill & Robertson 1966; Comeron et al. 2008). This occurs because drift randomly generates positive and negative associations between alleles with equal probability each generation, but positive associations are depleted more rapidly by selection than negative ones, since positive associations generate more genetic variation for selection to act on (Barton & Otto 2005). By analogy, selection on male trait and preference alleles may also interact with drift, potentially reshaping trait-preference correlations and the coevolutionary dynamics of male traits and female preferences. This possibility is particularly relevant given that median estimated effective population sizes are below 1,000 across taxonomic groups (Clarke et al. 2024).

How finite population size influences the coevolutionary dynamics of traits and preferences under Fisherian selection remains largely unknown, as nearly all previous models assume an infinite population size. Nichols and Butlin (1989) investigated the ecological constraints caused by finite population size and showed that mate choice can be less effective in smaller populations, since viability selection leaves only a small number of surviving males, limiting females’ mating opportunities. Although the authors also conjectured that genetic drift and recombination may further erode trait-preference correlations, the impacts of these genetic processes were not explicitly explored.

Here, I investigate how genetic drift and recombination rate influence the coevolution of male traits and female preferences via Fisherian sexual selection in finite populations. Specifically, I identify the conditions under which strong versus weak trait-preference genetic correlations arise, and determine whether the strength of these correlations reliably reflects the extent of coordinated evolution between male traits and female preferences.

## Methods

### Two-locus model

Following Kirkpatrick (1982), I consider a haploid polygynous population of constant population size with *N* females and *N* males. A locus T with alleles *T*_1_ and *T*_2_ governs a trait expressed only in males, and a locus P with alleles *P*_1_ and *P*_2_ determines the female mating preference. *T*_1_ males are without the secondary sexual characteristic, while *T*_2_ males bear a trait which makes them more attractive to *P*_2_ females than are *T*_1_ males. *P*_1_ females mate indiscriminately, while *P*_2_ females prefer to mate with a *T*_2_ male *a* times more frequently than with a *T*_1_ male. The male trait is subject to viability selection, and the fitness of *T*_1_ and *T*_2_ males is 1 and 1 + *s*, respectively. The selection coefficient *s* may be either positive or negative. I assume no direct selection at the P locus, so that there is no interaction between genetic drift and natural selection at the two loci, which could otherwise be confounding.

Each generation starts from the stage immediately after juveniles have been randomly sampled from the offspring produced by the previous generation. I assume that exactly *N* female and *N* male juveniles are sampled to avoid confounding effects of stochastic variation in sex ratio on altering effective population size and the strength of sexual selection. After the census, the population undergoes viability selection, mate choice and offspring production, where I assume the number of potential offspring is sufficiently large such that these processes can be treated deterministically. I denote the frequency of *T_i_* and *P_i_* (*i* = 1,2) in the juvenile population before selection by *t_i_* and *p_i_*, respectively. A common measure of genetic correlation is linkage disequilibrium (LD), *D* = *x_T_*_1_*_P_*_1_ *x_T_*_2_*_P_*_2_ − *x_T_*_1_*_P_*_2_ *x_T_*_2_*_P_*_1_, where *x_i_* is the frequency of haplotype *i*. However, since *D* also depends on the amount of genetic variation at the two loci, I additionally calculated the correlation coefficient, 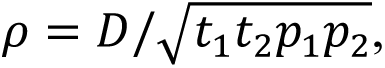 which provides a standardized measure of association by accounting for differences in genetic variation. This is also the metric reported in previous meta-analyses of trait-preference genetic correlations (Greenfield et al. 2014).

The two-locus system can be described by the frequency of male trait *T*_2_ (*t*_2_), frequency of preference allele *P*_2_ (*p*_2_) and LD. The deterministic recursions for *t*_2_, *p*_2_ and *D* in infinitely large populations are given by Equation (1) in Kirkpatrick (1982). In finite populations, allele frequency and LD are random variables, and our focus is their expectations, denoted by D[X]. Random sampling causes *t*_2_, *p*_2_ and *D* to differ from the values predicted by deterministic recursions, and we denote the perturbations that arise from a single generation of sampling by *ζ_t_* and *ζ_p_* and *ζ_D_*. The expectation of perturbations in allele frequencies and LD is 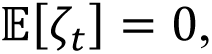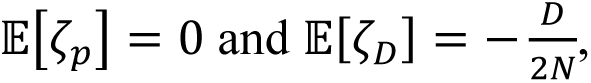, respectively (Barton & Otto 2005). The second moments of these perturbations is at the order of 1/*N* (Section 1 of SI Appendix). Perturbations from each generation will accumulate to cause the actual trajectory of *t*_2_, *p*_2_ and LD to deviate from the deterministic trajectory, and I denote the overall deviations in *t*_2_, *p*_2_ and LD by *δt*, *δp* and *δD*, respectively.

To analyze the source of the deviations and the relative contributions of perturbations in each dimension, I derive the recursions for the deviations, using the perturbation analysis approach of Barton & Otto (2005). This method assumes that the deviations remain small and the population size is large enough so that the male trait always goes to fixation. Briefly, denote *x* = (*t*_2_, *p*_2_, *D*), and let *x*^∗^ = *f*(*x*) be the deterministic recursions of allele frequencies and LD in infinitely large populations, given in Kirkpatrick (1982). The recursion of the deviation vector *z* = (*δt*, *δp*, *δD*) is *z*^∗^ = *f*(*x* + *z*) − *f*(*x*) + ***ζ***, where the vector ***ζ*** = (*ζ_t_*, *ζ_p_*, *ζ_D_*) represents the perturbations. The recursions of the first moments E[*δt*], E[*δp*] and E[*δD*] and second moments of deviations E[*δt*^2^], E[*δp*^2^], E[*δD*^2^], E[*δtδp*], E[*δtδD*] and E[*δpδD*] can be are given in Section 1 of the SI Appendix.

### Individual-based simulations for polygenic trait and preference

As an extension to the two-locus model, I also investigate the case when male traits and female preferences are polygenic by individual-based simulations (see Section 3 of the SI Appendix for detailed description; simulation code in C++ is available at XXX [archived upon acceptance]). Briefly, the simulation considers a diploid population with *N* females and *N* males. The quantitative male trait *z* and female preference *y* are controlled by *n_z_* and *n_y_* additive biallelic loci, respectively, each with identical allelic effects. Mutations occur at equal forward and backward rates at each locus, with total genomic mutation rates across all trait and preference loci being *U_z_* and *U_y_*, respectively. Male trait and preference loci are located along a chromosome, with an expected number of *L* crossover events during meiosis positions. Female mate choice follows an absolute preference function from Lande (1981). For a female with preference value *y*, the relative probability of mating with a male of trait value *z* is *ψ*(*z*|*y*) = *e*^−*ν*(*z*−*y*)2/2^, where *ν* controlls the strength of preference. Both the male trait and female preference are subject to stabilizing viability selection, and surviving females have equal fertility.

The relative viability of a male with trait value *z* is *w*(*z*) = *e*^−*ω*^*^zz^*^2/2^, where *ω* determines the strength of viability selection on the male trait. The relative fitness of a female with preference value *y*, arising from the costs of mate choice is *w*(*y*) = *e*^−*ω*^*^y^*^(*y*−*z*∗)2/2^, where *z*^∗^ is the mean male trait value after viability selection and *ω_y_* is the strength of selection acting on preference (Xu & Servedio 2026).

## Results

Similar to Servedio (2025), below I focus on two scenarios when both the male trait and preference loci are polymorphic, since otherwise there is no genetic correlation. The first scenario considers the sweeping phase when the allele frequency of male trait and preference alleles evolve toward 1 or 0 in finite populations, generating transient genetic correlations. The second scenario is when male trait and preference loci maintain polymorphism through a balance among mutation, selection, and genetic drift. For comparison, I first present the dynamics in infinitely large populations and examine whether key factors that increase trait-preference genetic correlations also promote trait-preference coevolution. I then show how results in finite populations qualitatively differ from the deterministic ones.

### Dynamics in infinitely large populations

Under Fisherian selection, the strength of sexual selection on the male trait depends on the frequency of preference allele, while selection on preference is proportional to both the strength of selection on the male trait and LD (Kirkpatrick 1982). When sexual selection outweighs the (potential) viability cost, the male trait sweeps to fixation, leading to an increase in preference frequency through LD built up over time (Figures 1a and 1b). In general, stronger trait-preference correlations often entail larger overall increases in preference frequency, although there are exceptions (the black and red curves exhibit concordant trends in Figures 1c-e but not in Figure 1f).

**Figure 1.**
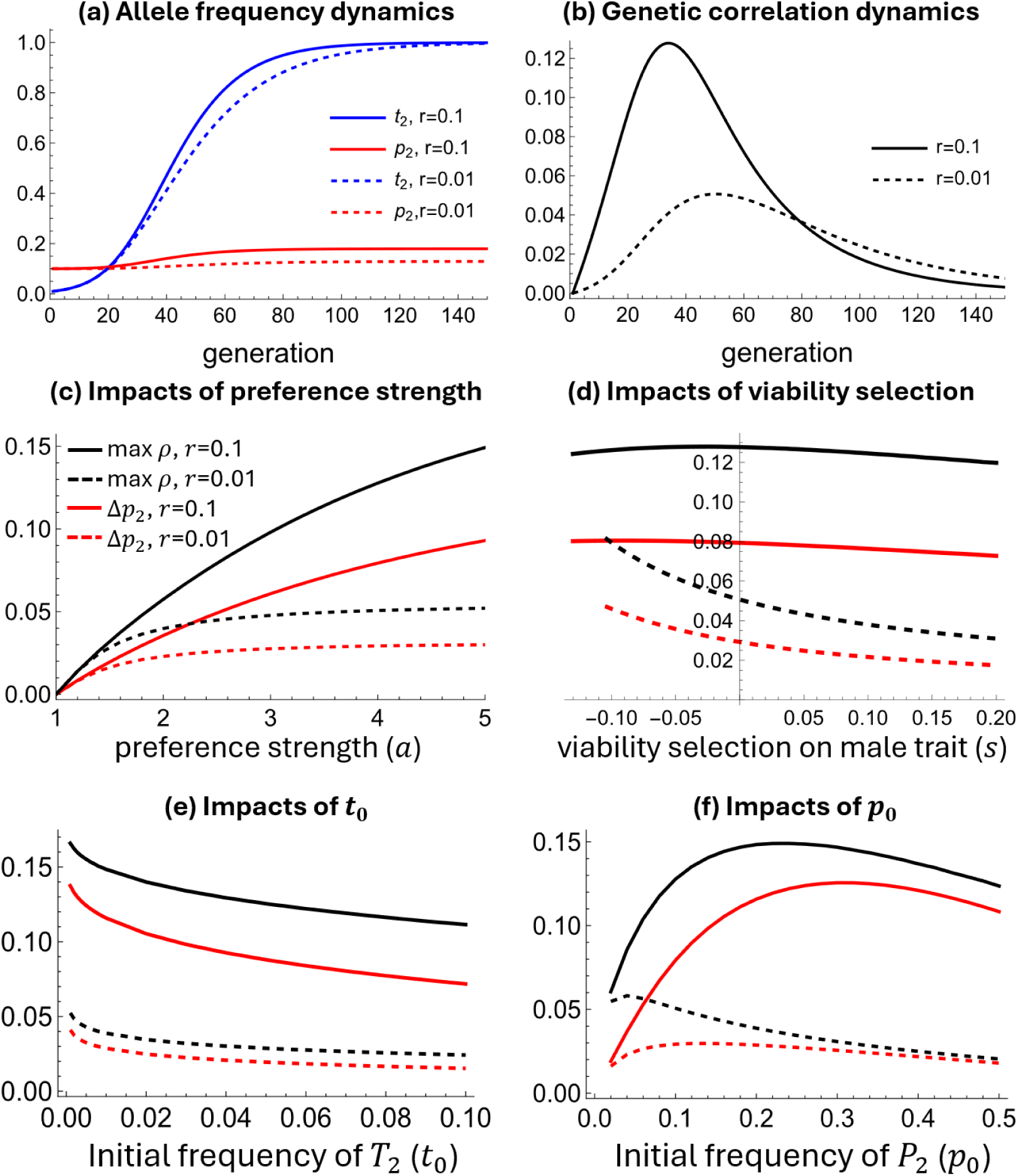
**(a) & (b):** Impacts of recombination rate on the Fisherian runaway dynamics in infinitely large populations. **(c)-(f):** Impacts of preference strength, viability selection and initial allele frequencies on the maximum trait-preference correlation coefficient during the sweep (max *ρ*), and the overall change in preference frequency when the male trait reaches fixation (Δ*p*). Unless otherwise specified, parameters are *s* = 0, *a* = 4, *t*_0_ = 0.01, *p*_0_ = 0.1, *D*_0_ = 0.

The strength of the trait-preference correlation is strongly influenced by the recombination rate between the trait and preference loci. Given that the trait and preference loci are initially at linkage equilibrium, more frequent recombination leads to a stronger correlation and thus a larger increase in preference frequency (compare the solid and dashed lines in Figures 1a-f), as also found in Servedio (2025). This is because at an early stage when the male trait is rare, mate choice makes recombination more effective at forming than breaking up *T*_2_*P*_2_ haplotype: *T*_2_*P*_2_ females mate disproportionately often with males carrying allele *T*_2_, and *T*_2_*P*_2_ males are more likely to be chosen by *P*_2_ females. Nevertheless, at a late stage when *T*_2_ males become common, more frequent recombination can lead to a faster decay of LD by breaking up *T*_2_*P*_2_ haplotype (compare the decline period of the solid and dashed curves in Figure 1b). In addition, when the trait and preference alleles are initially highly positively associated, more frequent recombination may instead promote the trait-preference correlation and preference evolution (Figure S1).

Trait-preference correlations and preference evolution also depend on the strength of sexual and viability selection on the male trait, and on the initial allele frequencies. Stronger preferences enhance genetic correlation by increasing the frequency of mating between *T*_1_*P*_2_ females and *T*_2_*P*_1_ males, and thus the generation of *T*_2_*P*_2_ haplotype via recombination. Therefore, this effect is pronounced only when the recombination rate is high (compare dashed and solid lines in Figure 1c). Reducing the relative viability or initial frequency of the male trait *T*_2_ promotes the trait-preference correlation as well as preference evolution (Figures 1d and 1e), because it slows the sweep of the male trait, allowing genetic correlations to accumulate over more generations.

However, the effect of the initial frequency of preference allele *P*_2_, *p*_0_, is non-monotonic, with the trait-preference correlation and the increase in preference frequency peaking at intermediate values of *p*_0_ (Figure 1f). Intuitively, the strength of sexual selection is weak when *p*_0_ is low, but when *p*_0_ is excessively high, the male trait sweeps too rapidly to allow genetic correlations to accumulate over generations. Importantly, increases in genetic correlation caused by changes in *p*_0_ may not correspond to greater evolution of preference frequency (compare the black and red lines in Figure 1f), because the evolution of preference depends on the cumulative product of LD and the strength of selection on the male, both of which are affected by *p*_0_.

### Dynamics in finite populations

Since evolutionary dynamics in finite populations are stochastic, below we focus on the expected dynamics averaged across replicates. Each generation, genetic drift generates perturbations in the frequencies of male trait and preference alleles, and in LD. Perturbations in any single dimension generate deviations from deterministic dynamics in all three dimensions in subsequent generations. Similar to the mechanism underlying the Hill-Robertson effect (Barton & Otto 2005), although positive and negative perturbations arise with equal likelihood, subsequent deviations decay at different rates, resulting in non-zero expected deviations. However, unlike the Hill-Robertson effect, in which the expected deviations in genetic correlation and allele frequencies are consistently negative, the dynamics are more complicated under Fisherian selection.

Under frequent recombination, the magnitude of deviations caused by drift is small, and the expected dynamics in finite populations closely follow the deterministic dynamics (Figures 1a-c). As recombination becomes less frequent, deviations generated by perturbations at each generation decay more slowly and accumulate, leading to larger magnitude. Therefore, the expected dynamics in finite populations can diverge substantially from deterministic ones (Figure 2c). The magnitude of deviations is also greater in smaller populations (Figure S2), and when the sweep of the male trait is slower — such as when its initial frequency is lower or when overall selection on the male trait is weaker owing to a weaker female preference or a stronger viability cost (Figure S3). This is because the variance of perturbations caused by drift is proportional to 1/*N*, and a slower sweep of the male trait allows perturbations to occur over more generations. Additionally, the male trait allele may be lost due to drift in finite populations, and among replicates where it ultimately fixes, deviations in both allele frequencies and LD are more positive than the average across all replicates (Figure S4).

**Figure 2.**
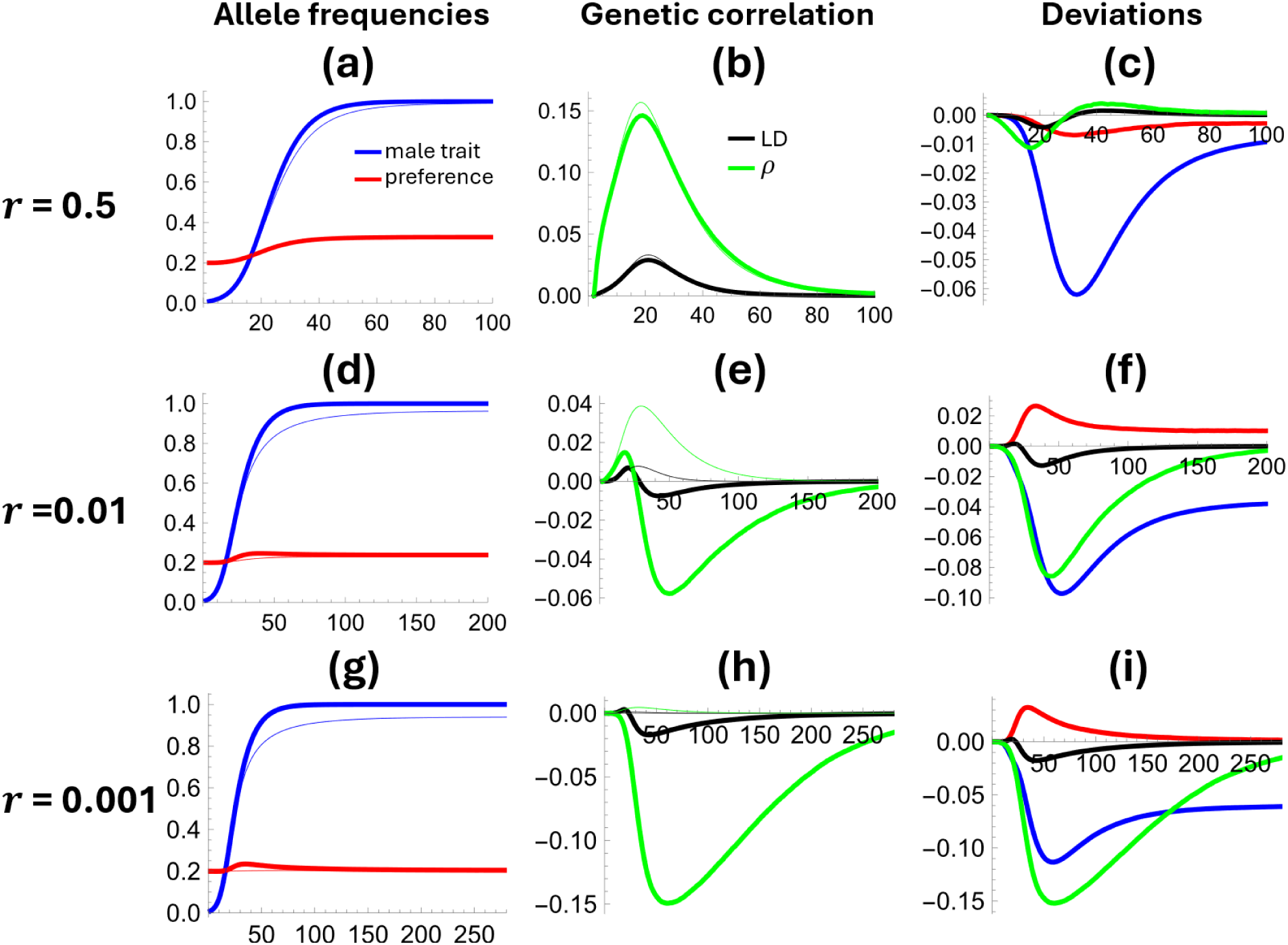
Expected dynamics of allele frequency and trait-preference correlations in finite populations and their deviations from the deterministic dynamics under different recombination rates *r*. Results are the average across 10^5^ simulation replicates. The first and second columns show dynamics of allele frequencies and genetic correlations (LD and correlation coefficient *ρ*) over generations, where the thick and thin lines are dynamics in finite and infinitely large populations, respectively. The third column shows the dynamics of deviations (differences between thick and thin lines in the first and second columns). Parameters: 2*N* = 1000, *a* = 4, *s* = 0, *t*_0_ = 0.01, *p*_0_ = 0.2, *D*_0_ = 0.

In general, genetic drift always slows the evolution of the male trait, producing negative expected deviations from deterministic dynamics (blue lines in Figure 2). Additional analysis shows that this is mainly driven by perturbations in the male trait and preference frequencies caused by drift (panels (a) and (d) in Figures S5 and S6; detailed in Section 2 of the SI Appendix).

In contrast, the expected deviations in preference frequency and trait-preference correlation depend on the recombination rate. When recombination is frequent, expected deviations in preference frequency remain negative (red line in Figure 1c), while the expected deviations in LD and the correlation coefficient are initially negative when the male trait *T*_2_ is rare, but become positive later as *T*_2_ becomes common (black and green lines in Figure 1c). When recombination is not too frequent, expected deviations in LD are initially slightly positive, and become negative later (black lines in Figures 2f and 2i); however, expected deviations in the correlation coefficient remain negative throughout (green lines in Figures 2f and 2i), suggesting that the positive deviations in LD at early stages are caused by changes in genetic variation. Although deviations in LD are largely negative, expected deviations in preference frequency remain positive (red lines in Figures 2f and 2i), meaning that drift promotes preference evolution.

Therefore, an important phenomenon is that although indirect selection on preference is proportional to LD within each replicate at every generation (Kirkpatrick 1982), when averaged across replicates, the expected genetic correlations may be decoupled from the expected changes in male trait and preference allele frequencies: even when the expected LD is negative (Figure 2e and 2h), the expected preference allele frequency can increase more than the deterministic prediction (red curves in Figures 2f and 2h).

This seemingly counterintuitive pattern is mainly driven by perturbations in LD generated by drift, although perturbations in allele frequencies also contribute (detailed in Section 2 of the Appendix). Specifically, during the early stage of Fisherian selection, LD builds up with accelerating returns (Figures 1a and 1b). Positive perturbations in LD generate even greater deviations in LD in subsequent generations, resulting in positive expected deviations in LD and thus also positive deviations in preference frequency (panels (b) and (c) in Figure S5). In the later stage when LD declines over time as the male trait approaches high frequency (Figure 1b), positive deviations in LD decay more rapidly than negative ones, resulting in negative expected deviations in LD (green line in Figure S6f). This is because positive perturbations in LD lead to faster-than-average evolution of preference, which generates stronger selection and thus faster depletion of genetic variation at the male trait locus. However, the increased selection on the male trait can increase indirect selection on preference (Kirkpatrick 1982), which leads to positive expected deviations in preference frequency (Figure S6e). In other words, although perturbations in LD by drift on average lead to negative deviations in LD, they still generate positive deviations in preference frequency by influencing male trait evolution.

### Average genetic correlation may not reflect trait-preference coevolution

In natural populations, different trait and preference loci may undergo selective sweeps at different times, so empirical estimates effectively average across loci under different stages of the sweep (Greenfield et al. 2014). If loci are approximately uniformly distributed across sweep stages, this cross-locus average approximates the temporal average over a single sweep. Therefore, I use the temporal average correlation coefficient and the overall allele-frequency changes during a sweep as summary measures of the strength of trait-preference association and the extent of evolution of the trait and preference. As shown below, across a wide range of recombination rates and population sizes, the average genetic correlation may not reflect the extent of trait-preference coevolution.

The temporal average of genetic correlations is positive only when recombination is sufficiently frequent (the region above white line in Figure 3a); otherwise, it becomes negative, but the male trait and preference alleles can still increase in frequency under these conditions (regions below the white line in Figures 3a-c).

**Figure 3.**
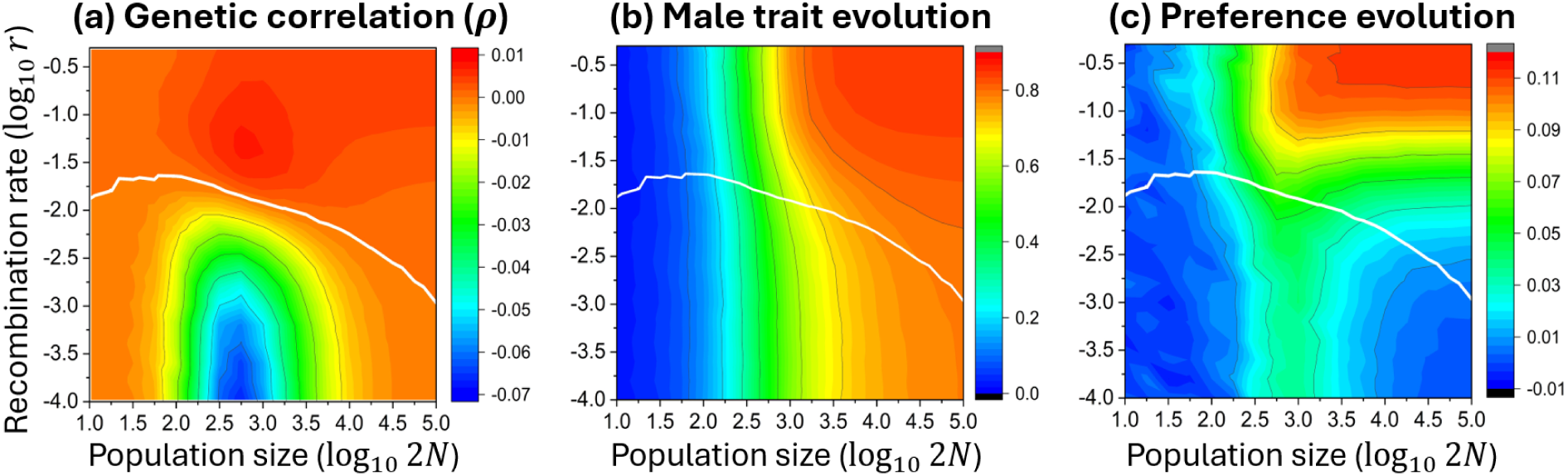
Impacts of recombination rates and population sizes on the trait-preference correlation and the evolution of male trait and preference allele. Panel (a) shows the trait-preference correlation coefficient *ρ* averaged over 200 generations. Panels (b) and (c) show changes in male trait frequency and preference allele frequency after 200 generations. Results are insensitive to the time length chosen. In all panels, the white line correspond to parameters when the average correlation coefficient is 0. Results are the average across 10^5^numerical simulation replicates. Parameters: *a* = 4, *s* = −0.1, *t*_0_ = 0.01, *p*_0_ = 0.2, *D*_0_ = 0.

When recombination is frequent, larger population sizes consistently promote the evolution of both male traits and preferences, primarily due to the higher fixation probability of the male trait (regions above the white line in Figures 3b and 3c), whereas the average genetic correlation peaks at intermediate population sizes (Figure 3a). When recombination is infrequent, intermediate population sizes generate the most negative genetic correlations, yet correspond to the greatest increase in preference frequency (compare the region around 2*N* ≈ 10^3^below the white line in Figures 3a and 3c). As shown in the previous section, under infrequent recombination, drift tend to generate negative expected deviations in genetic correlation and positive expected deviations in preference frequency (Figures 2f and 2i). The magnitude of these deviations is greatest at intermediate population sizes, since in large populations, drift is too weak to generate substantial deviations, while when population size is too small, the male trait is frequently lost (Figure 3b), so there is no genetic correlation. These patterns are qualitatively robust to different parameter values (Figure S7). Moreover, despite negative average genetic correlations, fixation of the male trait and preference alleles remains positively associated (Figure S8).

In addition to the temporal average, I also examine the maximum and minimum expected correlation coefficients attained during the sweeping phase. The magnitude of both statistics is greatest at intermediately small populations (Figure S9). However, the magnitude of neither metric reflects the extent of evolution of male traits and female preferences (compare Figure S9 with Figures 3b and 3c).

### Mutation-selection-drift balance

Previous sections focus on the sweeping phase of Fisher’s process, during which genetic correlations arise transiently. Here, I consider the case when male trait and preference loci maintain polymorphism under a balance among mutation, selection, and drift, generating persistent genetic correlations. Allele frequency and genetic correlation exhibit a probability distribution; below, I focus on their expected values. In general, provided that the population size is small enough, interactions between selection and drift tend to reduce trait-preference correlations (Figures 4a and 4d) while increasing the preference frequency compared to the deterministic prediction (Figures 4c and 4f). These patterns are qualitatively consistent with those observed during the sweeping phase and can be understood by referring to the mechanisms discussed in the previous section *Dynamics in finite populations*.

**Figure 4.**
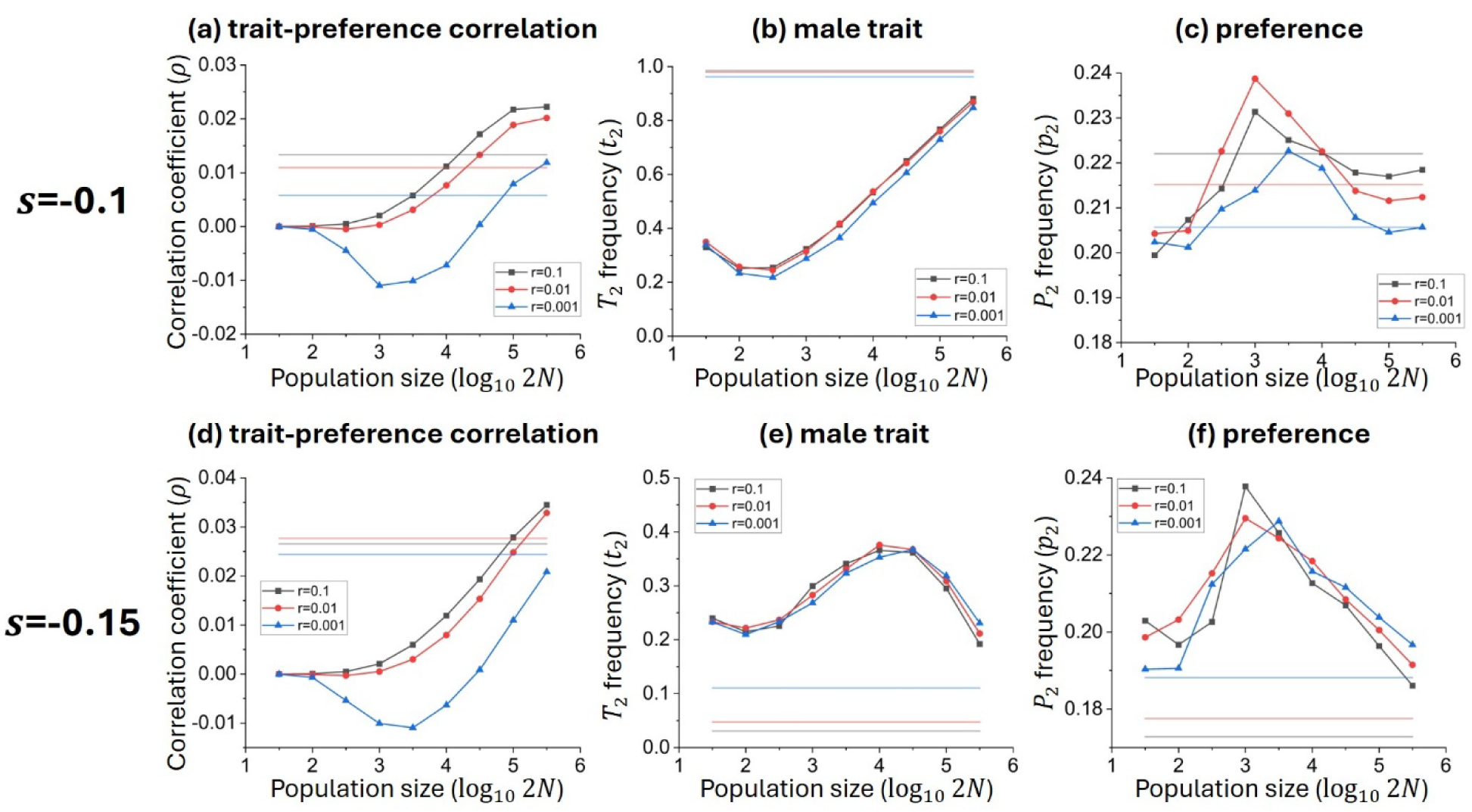
Expected allele frequencies and genetic correlations in finite populations when the male trait and preference loci maintain polymorphism under mutation-selection-drift balance. The upper and lower panels show cases when the male trait *T*_2_ is selected against and or favored in infinitely large populations, respectively, by changing viability selection *s*. Horizontal lines are deterministic predictions for infinitely large populations. Dots represent simulation results averaged across10^3^ replicates. Each replicate is first run for 12*N* generations to reach mutation-selection-drift balance, and due to temporal fluctuations, the statistics are calculated by averaging over an additional 6*N* generations. For the results presented, preference strength is *a* = 2. Mutation rates between alleles *T*_1_ and *T*_2_ at the male trait locus are symmetric at 0.0001. At the preference locus, the mutation rate from allele *P*_1_ to *P*_2_ is 0.00004, and 0.00016 from *P*_2_ to *P*_1_, so that *P*_2_ frequency is around 0.2.

However, under mutation-selection-drift balance, expected allele frequencies and genetic correlations are sensitive to even slight changes in selection strength, so drift can generate much larger deviations from deterministic predictions than in the sweeping scenario. This leads to several differences. Specifically, when the male trait allele would be selected against and maintained at low frequency in infinitely large populations, it can instead be favored and maintained at a much higher frequency in finite populations (Figure 4e), because drift elevates preference frequency (Figure 4d) and thereby strengthens sexual selection on the male trait. Moreover, when population size is sufficiently large, the expected trait-preference correlation may exceed deterministic predictions (Figures 4a and 4d). This occurs because genetic drift increases preference allele frequency (Figures 4c and 4f) and increases genetic variation at the male trait locus (male trait frequency is closer to 0.5 in Figures 4b and 4e).

The above results suggest that when male trait and preference are controlled by multiple loci subject to selection of varying direction and magnitude, positive and negative genetic correlations among pairwise trait and preference loci may offset one another, resulting in weak overall genetic correlations. Consistent with this proposition, individual-based simulations show that the expected genetic correlations between polygenic male traits and preferences tend to be weaker in smaller populations and when recombination is less frequent, even when genetic variation of male traits and preference is controlled at similar levels across population sizes and recombination rates (Figure 5a). When genetic variance is allowed to coevolve, the reduction in genetic correlations is even greater, owing to lower levels of genetic variation in both male traits and preferences in smaller populations and under tighter linkage (Figure S11) (Hall et al. 2000; Xu & Servedio 2026).

**Figure 5.**
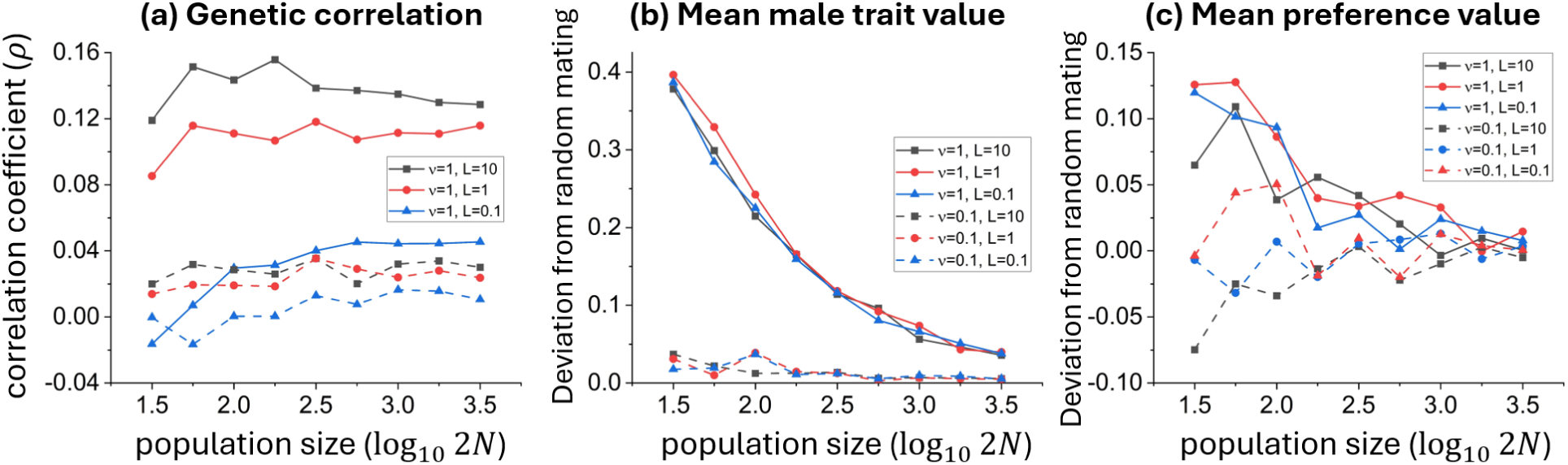
Effects of population size and recombination frequency on trait-preference genetic correlations (a) and on the coevolution of mean male trait and preference values (b) and (c), when male traits and female preferences are polygenic and maintained under mutation-selection-drift balance. Results are from individual-based simulations, averaged across 20,000 generations after the system reaches mutation-selection-drift balance (see *Materials and Methods*), with both male traits and preferences controlled by 20 additive loci. Females exhibit absolute preferences following Lande (1981), with preference strength determined by *νν*. Solid and dashed lines show results under strong (*νν* = 1) and moderate (*νν* = 0.1) preferences, respectively. Panels (b) and (c) show the deviation of the mean male trait value from its viability optimum relative to that in a randomly mating population. Different colors represent different expected numbers of crossovers (*L*) occurring among all loci. Genetic variation in male traits and preferences is controlled at similar levels across population sizes and recombination rates by adjusting allele effect sizes (Figure S10), thereby eliminating the confounding effects of genetic variance on genetic correlations. Other parameter values are: genetic variance of the male trait and preference when all loci have frequency 0.5 at linkage equilibrium, *G_x_* = *G_y_* = 1; genomic mutation rates for the male trait and preference, *U_z_* = *U_y_* = 0.1; and strengths of stabilizing viability selection on the male trait and preference, *ω_z_* = 0.1 and *ω_y_* = 0.02, respectively.

Despite the reduction in expected trait-preference genetic correlations, the coevolution of mean male trait and female preference values may be enhanced. The evolution of mean trait values can proceed in either positive or negative directions depending on the initial state (Lande 1981), and small populations experience greater fluctuations. To account for this, I calculate the temporal average deviation of male trait values and female preference values from their respective viability optima, relative to those in randomly mating populations, thereby eliminating fluctuations in trait means caused by drift alone. Although the trait-preference correlation is weaker in smaller populations (Figure 5a), the extent of evolution of both male traits and female preferences becomes larger, particularly when preference strength is strong (Figures 5b and 5c). Also, although lower recombination frequencies substantially reduce trait-preference correlations (compare the lines for *L*=0.1, 1, 10 in Figure 5a), they do not translate into a smaller extent of evolution of male traits and female preferences (compare the lines for *L*=0.1, 1, and 10 in Figures 5b and 5c). Together, these results further suggest that the strength of genetic correlations may be largely decoupled from the extent of coevolution between male traits and female preferences in natural populations.

## Discussion

Fisherian sexual selection is a process driven by the buildup of genetic correlations between male display traits and female preferences (Lande 1981; Kirkpatrick 1982; Mead & Arnold 2004). Correspondingly, the strength of such correlations has long been considered a key indicator of the effectiveness of Fisher’s process in driving the coevolution of exaggerated display traits and preferences. However, empirical studies often fail to detect such correlations even with substantial variation in both traits and preferences (Ritchie et al. 2005; Zhou et al. 2011; Greenfield et al. 2014), leaving the prevalence of Fisher’s process in nature equivocal. This study suggests a potential resolution to this dilemma. I show that Fisher’s process in finite populations is not simply a noisy version of the deterministic expectation; instead, interactions between Fisherian selection and genetic drift can fundamentally decouple expected trait-preference genetic correlations from changes in male trait and female preference allele frequencies. In natural populations, genetic correlations averaged across loci may often be weak or even negative, yet substantial coevolution of male trait and preference alleles may still occur, given that median estimated effective population sizes of various taxonomic groups are below 1,000 (Clarke et al. 2024).

The key underlying mechanism is that interactions between Fisherian selection and genetic drift tend to reduce expected trait-preference genetic correlations while increasing the expected change in female preference frequency beyond deterministic expectations (Figures 2 and 4), although within each replicate, selection strength on preference remains proportional to LD. These deviations arise because drift generates random perturbations in allele frequencies and LD each generation, which influence subsequent dynamics and cause further departures from deterministic trajectories. However, deviations resulting from positive and negative perturbations do not decay symmetrically, leading to non-zero expected deviations when averaged across replicates (Section 2 of the SI Appendix).

Briefly, when the male trait is rare and LD builds up via positive feedback (Figures 1a and 1b), positive perturbations in LD amplify more strongly than negative ones, generating a positive net deviation in LD and, consequently, in preference frequency. As the male trait becomes common and LD decays, positive perturbations dissipate more rapidly than negative ones, generating a negative net deviation in LD, analogous to the classical Hill-Robertson effect described in the introduction (Barton & Otto 2005). Preference frequency nonetheless retains a positive net deviation throughout, because indirect selection on preference depends not only on LD but also on the strength of selection on the male trait (Kirkpatrick 1982): positive LD perturbations accelerate preference evolution, which intensifies selection on the trait and further boosts indirect selection on preference, offsetting the negative LD bias.

More generally, the above mechanisms can be viewed as an extension of the classical Hill-Robertson effect, which may extend well beyond sexual selection to any evolutionary system involving positive feedback between interacting loci. For example, when two beneficial alleles exhibit positive epistasis such that selection generates positive LD between them, and the strength of selection on one allele increases with LD, deviations from deterministic dynamics are qualitatively similar to those observed under Fisherian sexual selection (Figure S12). Therefore, although the current model assumes females either mate randomly or express a preference, similar dynamics are expected whenever the coevolution of traits and preferences involves self-reinforcing feedback, including cases where females consistently prefer a particular male trait but vary in preference strength (Kirkpatrick 1982; Xu et al. 2023), or where females carrying different preference alleles prefer different male traits (Servedio & Bürger 2018).

### Implications for empirical tests of Fisherian sexual selection

As Greenfield et al. (2014) pointed out, observed trait-preference genetic correlations are expected to represent the combined contributions of trait and preference loci in different evolutionary states. Individual loci may be fixed, maintained under stable polymorphisms, or undergoing selective sweeps (Tinghitella et al. 2018; Gallagher et al. 2022). Despite these differences, our results reveal a qualitatively consistent impact of genetic drift across these scenarios; instead, population size and recombination frequency among trait and preference loci are key determinants of the extent to which genetic drift reduces trait-preference genetic correlations and promotes the evolution of preference.

Therefore, whether drift plays an important role in trait-preference coevolution depends on the genetic architecture of display traits and preferences, which can vary with traits and species. In some species, traits and preferences are mainly contributed by few large-effect loci located on different chromosomes or linkage groups (Limousin et al. 2012; Blankers et al. 2019). In this case, the impacts of drift may be unimportant. On the other hand, genetic analyses have revealed extensive co-localization and tight physical linkage between major-effect loci underlying mating signals and preferences in *Laupala* crickets, *Heliconius* butterflies, and *Drosophila* (McNiven & Moehring 2013; Byers et al. 2021; Xu & Shaw 2021; Ritchie & Butlin 2024; VanKuren et al. 2025; Xu & Shaw 2026). Such genetic architectures might contribute to the weak or slightly negative trait-preference genetic correlations reported in other species from the same taxonomic groups (Greenfield et al. 2014).

Empirical studies measuring trait-preference genetic correlations have rarely reported population sizes (Greenfield et al. 2014), so our prediction that genetic correlations should generally be weaker in smaller populations warrants further empirical investigation. This prediction could be tested experimentally by manipulating population size in laboratory populations. However, our individual-based simulations find large stochastic fluctuations in genetic correlations in small populations, which can introduce substantial noise into empirical estimates. This may explain why trait-preference genetic correlations were significantly positive in field populations of *Gryllus texensis* but negative in the subsequent laboratory generation (Gray & Cade 1999a, 1999b). Therefore, reliable estimation of trait-preference genetic correlations may require measurements across multiple replicate populations and multiple generations. Furthermore, evolution of display traits and preferences in small populations may not necessarily involve strong genetic correlation. As indirect selection on preferences tends to be weak (Servedio 2025), preference evolution may be dominated by genetic drift in small populations. Drift can randomly elevate preference allele frequencies or the mean preference value, and concurrently reduces genetic variation in preference, which allow exaggerated display traits to evolve subsequently without strong concomitant trait-preference genetic correlations (Henshaw et al., 2022). This scenario is particularly plausible when preference architectures are governed by few major-effect loci.

Individual trait-preference locus pairs may also contribute unequally, and sometimes in opposite directions, to the overall genetic correlation, since loci underlying male traits and female preferences can differ substantially in their effect sizes and recombination rates (Wright et al. 2008; Chenoweth & McGuigan 2010; Limousin et al. 2012; Xu & Shaw 2026). Therefore, rather than focusing on the overall genetic correlation, investigating how genetic association varies across individual trait-preference locus pairs, depending on their effect sizes and recombination rates, may provide a more powerful test of the current theoretical predictions and a deeper understanding of sexual selection in nature.

### Future directions

Our two-locus models assume no direct selection on preferences, to eliminate the added complexity caused by the Hill-Robertson effect that would arise if there were direct selection acting on both the male trait and preference loci simultaneously. However, our individual-based simulations suggest adding direct selection on preferences may not qualitatively alter the core conclusions. Selection acting on preferences can still produce subtle nuanced effects. For instance, genetic drift generates negative association between two beneficial alleles and thereby effectively weakens their selective advantage (Barton & Otto 2005). When preferences incur survival costs while male display traits are adaptive and under positive selection, drift may strengthen trait–preference genetic correlations and mitigate selection against mating preferences. Furthermore, the handicap principle represents another major mechanism of sexual selection: male ornamental traits function as honest signals of underlying genetic quality (Iwasa et al. 1991; Dhole et al. 2018). Under this framework, the interplay between genetic drift, viability selection, and sexual selection targeting preference, display, and fitness loci creates far more complex evolutionary dynamics. Future work integrating direct selection on female mating preferences alongside handicap trait scenarios will therefore be valuable for fully unpacking natural sexual selection dynamics.

Fisher’s process is not only central to sexual selection theory, but also plays an important role in broader evolutionary processes, including adaptation (Kokko & Brooks 2003; Candolin & Heuschele 2008) and speciation (Servedio & Noor 2003; Ritchie et al. 2007; Servedio & Boughman 2017; Ritchie & Butlin 2024). Most theoretical studies that invoke Fisherian sexual selection as a mechanism driving speciation have assumed infinitely large populations. Our results suggest that relaxing this assumption may reveal evolutionary dynamics that are absent from deterministic models. Explicit consideration of the effects of finite population size may represent a fruitful direction for future sexual selection research.

## Acknowledgements

We are very grateful for useful comments from Matthew Osmond, Puneeth Deraje, Mete Yuksel, Jonathan Henshaw and, especially, Maria Servedio.

## References

1. Bailey, N. W., & Moore, A. J. (2012). Runaway sexual selection without genetic correlations: social environments and flexible mate choice initiate and enhance the Fisher process. Evolution, 66(9), 2674–2684.

2. Barton, N. H., & Otto, S. P. (2005). Evolution of recombination due to random drift. Genetics, 169(4), 2353–2370.

3. Blankers T., Berdan, E. L., Hennig, R. M., & Mayer, F. (2019). Physical linkage and mate preference generate linkage disequilibrium for behavioral isolation in two parapatric crickets. Evolution, 73(4), 777–791.

4. Bulmer, M. (1989). Structural instability of models of sexual selection. Theoretical population biology, 35(2), 195–206.

5. Byers, K. J., Darragh, K., Fernanda Garza, S., Abondano Almeida, D., Warren, I. A., Rastas, P. M.,…& Jiggins, C. D. (2021). Clustering of loci controlling species differences in male chemical bouquets of sympatric Heliconius butterflies. Ecology and evolution, 11(1), 89–107.

6. Candolin, U., & Heuschele, J. (2008). Is sexual selection beneficial during adaptation to environmental change?. Trends in ecology & evolution, 23(8), 446–452.

7. Chaine, A. S., & Lyon, B. E. (2008). Adaptive plasticity in female mate choice dampens sexual selection on male ornaments in the lark bunting. Science, 319(5862), 459–462.

8. Chenoweth, S. F., & McGuigan, K. (2010). The genetic basis of sexually selected variation. Annual Review of Ecology, Evolution, and Systematics, 41(1), 81–101.

9. Clarke, S. H., Lawrence, E. R., Matte, J. M., Gallagher, B. K., Salisbury, S. J., Michaelides, S. N., Koumrouyan, R., Ruzzante, D. E., Grant, J. W. & Fraser, D. J. (2024). Global assessment of effective population sizes: Consistent taxonomic differences in meeting the 50/500 rule. Molecular Ecology, 33(11), e17353.

10. Comeron, J. M., Williford, A., & Kliman, R. M. (2008). The Hill–Robertson effect: evolutionary consequences of weak selection and linkage in finite populations. Heredity, 100(1), 19–31.

11. Darwin, C. The Descent of Man and Selection in Relation to Sex (Charles Murray, London, UK, 1871).

12. Dhole, S., Stern, C. A., & Servedio, M. R. (2018). Direct detection of male quality can facilitate the evolution of female choosiness and indicators of good genes: Evolution across a continuum of indicator mechanisms. Evolution, 72(4), 770–784.

13. Fisher, R. A. The Genetical Theory of Natural Selection (Clarendon, Oxford, 1930).

14. Fry, J. D. (2022). A reexamination of theoretical arguments that indirect selection on mate preference is likely to be weaker than direct selection. Evolution Letters, 6(2), 110–117.

15. Gallagher, J. H., Zonana, D. M., Broder, E. D., Herner, B. K., & Tinghitella, R. M. (2022). Decoupling of sexual signals and their underlying morphology facilitates rapid phenotypic diversification. Evolution Letters, 6(6), 474–489.

16. Gray, D. A., and W. H. Cade. 1999a. Quantitative genetics of sexual selection in the field cricket, Gryllus integer. Evolution, 53(3), 848–854.

17. Gray, D. A., & Cade, W. H. (1999). Correlated-response-to-selection experiments designed to test for a genetic correlation between female preferences and male traits yield biased results. Animal behaviour, 58(6), 1325–1327.

18. Greenfield, M. D., & Rodriguez, R. L. (2004). Genotype–environment interaction and the reliability of mating signals. Animal Behaviour, 68(6), 1461–1468.

19. Greenfield, M. D., Alem, S., Limousin, D., & Bailey, N. W. (2014). The dilemma of Fisherian sexual selection: mate choice for indirect benefits despite rarity and overall weakness of trait-preference genetic correlation. Evolution, 68(12), 3524–3536.

20. Hall, D. W., Kirkpatrick, M., & West, B. (2000). Runaway sexual selection when female preferences are directly selected. Evolution, 54(6), 1862–1869.

21. Henshaw J. M., Fromhage, L., & Jones, A. G. (2022). The evolution of mating preferences for genetic attractiveness and quality in the presence of sensory bias. Proceedings of the National Academy of Sciences, 119(33), e2206262119.

22. Hill, W. G., & Robertson, A. (1966). The effect of linkage on limits to artificial selection. Genetics Research, 8(3), 269–294.

23. Iwasa, Y., Pomiankowski, A., & Nee, S. (1991). The evolution of costly mate preferences II. The “handicap” principle. Evolution, 45(6), 1431–1442.

24. Kirkpatrick, M. (1982). Sexual selection and the evolution of female choice. Evolution, 36(1), 1–12.

25. Kirkpatrick, M., & Barton, N. H. (1997). The strength of indirect selection on female mating preferences. Proceedings of the National Academy of Sciences, 94(4), 1282–1286.

26. Kokko, H., & Brooks, R. (2003). Sexy to die for? Sexual selection and the risk of extinction. In Annales Zoologici Fennici (pp. 207–219). Finnish Zoological and Botanical Publishing Board.

27. Kokko, H., & Heubel, K. (2008). Condition-dependence, genotype-by-environment interactions and the lek paradox. Genetica, 134(1), 55–62.

28. Kuijper, B., Pen, I., & Weissing, F. J. (2012). A guide to sexual selection theory. Annual Review of Ecology, Evolution, and Systematics, 43(1), 287–311.

29. Lande, R. (1981). Models of speciation by sexual selection on polygenic traits. Proceedings of the National Academy of Sciences, 78(6), 3721–3725.

30. Limousin, D., Streiff, R., Courtois, B., Dupuy, V., Alem, S., Greenfield, M. D. (2012) Genetic Architecture of Sexual Selection: QTL Mapping of Male Song and Female Receiver Traits in an Acoustic Moth. PLoS One. 7(9): e44554.

31. McNiven, V. T., & Moehring, A. J. (2013). Identification of genetically linked female preference and male trait. Evolution, 67(8), 2155–2165.

32. Mead, L. S., & Arnold, S. J. (2004). Quantitative genetic models of sexual selection. Trends in Ecology & Evolution, 19(5), 264–271.

33. Nichols, R. A., & Butlin, R. K. (1989). Does runaway sexual selection work in finite populations?. Journal of Evolutionary Biology, 2(4), 299–313.

34. Prum, R. O. (2010). The Lande–Kirkpatrick mechanism is the null model of evolution by intersexual selection: implications for meaning, honesty, and design in intersexual signals. Evolution, 64(11), 3085–3100.

35. Ritchie, M. G. (2007). Sexual selection and speciation. Annual Review of Ecology, Evolution, and Systematics, 38(1), 79–102.

36. Ritchie, M. G., & Butlin, R. K. (2024). Genetic coupling of mate recognition systems in the genomic era. Cold Spring Harbor Perspectives in Biology, 16(4), a041437.

37. Ritchie, M. G., Saarikettu, M., & Hoikkala, A. (2005). Variation, but no covariance, in female preference functions and male song in a natural population of Drosophila montana. Animal Behaviour, 70(4), 849–854.

38. Servedio, M. R. (2025). The contributions of direct and indirect selection to the evolution of mating preferences. Evolution, 79(1), 51–64.

39. Servedio, M. R., & Boughman, J. W. (2017). The role of sexual selection in local adaptation and speciation. Annual Review of Ecology, Evolution, and Systematics, 48, 85–109.

40. Servedio, M. R., & Noor, M. A. (2003). The role of reinforcement in speciation: theory and data. Annual review of ecology, evolution, and systematics, 34(1), 339–364.

41. Servedio, M. R., & Bürger, R. (2018). The effects on parapatric divergence of linkage between preference and trait loci versus pleiotropy. Genes, 9(4), 217.

42. Tinghitella, R. M., Broder, E. D., Gurule-Small, G. A., Hallagan, C. J., & Wilson, J. D. (2018). Purring crickets: the evolution of a novel sexual signal. The American Naturalist, 192(6), 773–782.

43. VanKuren N. W., Buerkle N. P., Lu W., Westerman E. L., Im A. K., Massardo, D., Southcott, L., Palmer, S. E., & Kronforst, M. R. (2025). Genetic, developmental, and neural changes underlying the evolution of butterfly mate preference. PLOS Biology, 23(3), e3002989.

44. Wright, D., Kerje, S., Brändström, H., Schütz, K., Kindmark, A., Andersson, L., Jensen, P., & Pizzari, T. (2008). The genetic architecture of a female sexual ornament. Evolution, 62(1), 86–98.

45. Xu, K., & Servedio, M. R. (2026). When sexual selection through mate choice depletes versus exaggerates genetic variation: Unraveling the lek paradox. Proceedings of the National Academy of Sciences of the United States of America, 123(16).

46. Xu, K., Lerch, B. A., & Servedio, M. R. (2023). The Fisher process of sexual selection with the coevolution of preference strength. Evolution, 77(4), 1043–1055.

47. Xu, M., & Shaw, K. L. (2021). Extensive linkage and genetic coupling of song and preference loci underlying rapid speciation in Laupala crickets. Journal of Heredity, 112(2), 204–213.

48. Xu, M., & Shaw, K. L. (2026). Linked song and preference loci suggest substantial contribution of genetic coupling in the rapid speciation of the Laupala crickets. Genetics, 233(1), iyag073.

49. Zhou, Y., Kelly, J. K., & Greenfield, M. D. (2011). Testing the fisherian mechanism: examining the genetic correlation between male song and female response in waxmoths. Evolutionary Ecology, 25(2), 307–329.

